# How Cancer Genomics Drives Cancer Biology: Does Synthetic Lethality Explain Mutually Exclusive Oncogenic Mutations?

**DOI:** 10.1101/091173

**Authors:** Harold Varmus, Arun M. Unni, William W. Lockwood

## Abstract

Large-scale analyses of cancer genomes are revealing patterns of mutations that suggest biologically significant ideas about many aspects of cancer, including carcinogenesis, classification, and preventive and therapeutic strategies. Among those patterns is “mutual exclusivity”, a phenomenon observed when two or more mutations that are commonly observed in samples of a type of cancer are not found combined in individual tumors. We have been studying a striking example of mutual exclusivity: the absence of co-existing mutations in the *KRAS* and *EGFR* proto-oncogenes in human lung adenocarcinomas, despite the high individual frequencies of such mutations in this common type of cancer. Multiple lines of evidence suggest that toxic effects of the joint expression of *KRAS* and *EGFR* mutant oncogenes, rather than loss of any selective advantages conferred by a second oncogene that operates through the same signaling pathway, are responsible for the observed mutational pattern. We discuss the potential for understanding the physiological basis of such toxicity, for exploiting it therapeutically, and for extending the studies to other examples of mutual exclusivity.

## Introduction

For more than three decades, the study of carcinogenesis has focused increasingly on the mutations that drive the conversion of cells that populate normal tissues into cells that display the hallmarks of cancer: excessive growth, an ability to invade, a failure to die or differentiate, and many other features of malignancy (Hanahan and Weinberg 2000; Hanahan and Weinberg 2011). Such mutations may arise stochastically as a result of errors in DNA replication or cell division, or they may be attributable to environmental, genetic, or behavior factors that promote mutations by damaging DNA directly or by impeding DNA repair (Stratton et al. 2009). Databases cataloging these mutations have been expanding exponentially under the influence of much more rapid methods for DNA sequencing and other modes of characterization and with investments in large scale cancer genome projects, such as such as The Cancer Genome Atlas (TCGA; https://cancergenome.nih.gov/) or the International Consortium for the Genomics of Cancer (ICGC; http://icgc.org/).

*General features of oncogenic mutations*. Individual somatic mutations identified in human cancers may be classified based on the chemical and physical changes in genetic material (base substitutions; small or large deletions or insertions; rearrangements such as DNA amplifications or chromosomal translocations) or on the functional consequences of the mutations (gain or loss of function)(Futreal et al. 2004; Watson et al. 2013). Gain-of-function (GOF) mutations that convert protooncogenes into active oncogenes often result from specific base substitutions, chromosomal translocations, or gene amplifications that strengthen gene function by increasing the abundance, changing the catalytic properties, or eliminating regulatory control of the gene product. Loss-of-function (LOF) mutations, such as inactivating missense or nonsense mutations or deletions, often combined with loss of heterozygosity, usually by intrachromosomal deletion or chromosomal loss, affect tumor suppressor genes (TSGs); similar functional effects can be achieved by selected nucleotide substitutions that confer a “dominant negative” effect. It is generally assumed that such LOF and GOF mutations offer a selective advantage, such as faster growth or less frequent cell death. When they occur in a cancer’s founding cell, they will be present in every cell in the cancerous clone; when they occur subsequently, they will be found in all cells in an evolutionary “branch” or subclone of the tumor.

Many oncogenic mutations are associated with certain types of cancers at only relatively low frequencies, but some mutations-such as GOF alleles of mutant *KRAS* in pancreatic adenocarcinomas, LOF mutations in the *RB* gene in small cell lung cancers, and the Philadelphia chromosome, harboring the BCR-ABL oncogene in chronic myeloid leukemia---are highly associated with a pathological diagnosis. Such findings suggest that cells in different tissues are susceptible to the cancer-inducing effects of specific mutations, a proposal that needs validation by experiments that expose cells at different stages of development in different organs to various mutations associated with established cancers.

Other mutations may occur coincidentally in the founding clone, without making a functional contribution to the cancer phenotype; those are considered “passenger” mutations and are rarely found recurrently in multiple tumors. Other biochemical or “epigenetic” changes in chromosomes (e.g. altered patterns of DNA methylation or modification of chromatin proteins) are also common in cancers (Meyerson et al. 2010), as are alterations in gene expression in adjacent, non-cancerous cells that constitute a tumor’s microenvironment; but the potentially important roles of the epigenome (Baylin and Jones 2016) and the microenvironment (Hanahan and Coussens 2012) in the initiation and maintenance of the oncogenic state have not been clearly defined.

*Genomic complexities*. Even when an analysis is confined to changes in the cancer cell’s DNA sequence (genotype), without an effort to discern functional consequences, the situation is complicated by two fundamental features: tumors are usually evident only after multiple mutations have occurred, and many cells in a tumor undergo subsequent mutations that contribute to the evolution of the cancer. These features mean that cancer genomes are complex, unique, internally heterogeneous, and changeable over time.

Such complexities are daunting. Still, recognizable genotypic patterns have emerged as new methods in genomics, such as “next generation sequencing” (NGS), have been applied to many tumor samples from a wide variety of histopathological types (Garraway and Lander 2013). As a result, both the individual mutations and combinations of lesions found at significant frequency in various histological types of cancers are increasingly used as criteria for making accurate diagnoses, which inform therapeutic decisions and prognostic advice (Garraway 2013). Further, analysis of the genetic alterations in a tumor (the “mutational signature”) is a useful guide to exogenous and endogenous factors that contribute to tumorigenesis by chemically damaging or misreplicating DNA (Lawrence et al. 2013). Even mutant alleles present at low frequencies in a tumor or mutations that appear during tumor growth may be useful clinically as indicators of subclones resistant to “targeted” drugs or endowed with augmented pathogenic potential.

*Mutational patterns and mutual exclusivity*. In addition to associating individual mutations with certain types of cancer and cancer phenotypes, the genotyping of many cancers has identified patterns of apparently cooperating mutations that are provocative but poorly understand. One such pattern is the co-appearance of certain mutations in certain tumor types---such as the dual LOF mutations in *TP53* and *RB* genes in small cell lung cancers or the constellation of mutations that promote Wnt signaling (LOF mutations of *APC* or GOF mutations in beta-catenin, R-spondin, or Axin), inactivate *TP53*, and activate *KRAS* in colorectal cancers (Cancer Genome Atlas 2012; George et al. 2015).

Another striking pattern---to be discussed at greater length here---is the absence of certain pairs of mutations that are individually frequent in certain tumor types. This pattern of “mutual exclusivity” is most easily recognized by displaying the well-recognized, recurrent mutations in a set of one type of tumors, using software such as that employed at the cBioPortal website (http://www.cbioportal.org) (Cerami et al. 2012). We have used statistical methods to determine the likelihood of such mutual exclusivities and judge their functional significance (Figure 1).

**Fig. 1.**
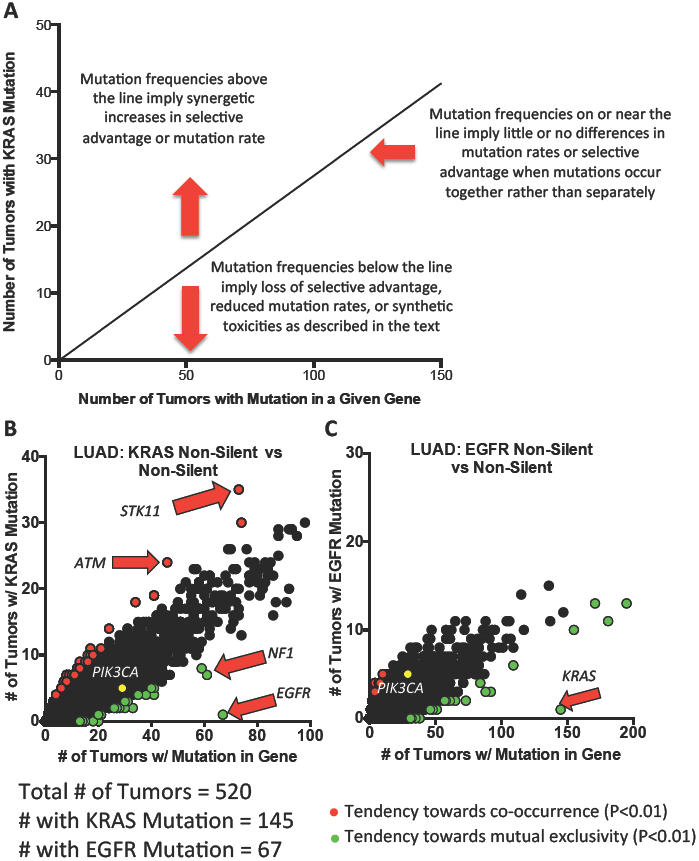
Assessing mutational relationships in lung adenocarcinoma. **A)** The diagram demonstrates the method used to identify gene pairs in which mutations are negatively associated (mutually exclusive) in cancer types, as well as those that are positively associated and those for which the frequency of detection is not influenced by co-existence in the tumor. With the assumptions regarding mutational frequencies and selective advantages or disadvantages, as described in the text, the method is based on the prediction that the more frequently any gene is mutated in a population of lung adenocarcinomas (x-axis), the greater the probability that it will occur in a tumor with a mutation in *KRAS* (y-axis). 2X2 Fisher’s Exact Tests are then used to determine non-random associations as previously described (Unni et al. 2015). **B and C)** In these panels, each point represents a unique gene, but data points overlap when multiple genes are mutated in the same number of tumors. Red dots represent genes for which non-silent mutations are positively associated with proto-oncogene mutations in the tumors (Right side Fisher’s Exact Test P≤0.01); green dots represent genes for which mutations are negatively associated with mutant proto-oncogenes (Left side Fisher’s Exact Test P<0.01). Yellow dots represent some important genes that are co-mutated at the “expected” rate. Genes of special interest are indicated by the arrows. **B)** Co-occurrence of non-silent mutations in other genes with missense *KRAS* mutations in lung adenocarcinoma (LUAD). The frequency may be changed from the expected number by differences in mutations rates or by changes in the selective advantages or disadvantages of the second mutation occurring in the presence of the first. Those changes may include toxic effects on the cell in which the second mutation occurs, as discussed extensively in the text. These factors will account for genes which are detected in mutant forms more or less frequently than expected with mutant *KRAS*. These include *STK11* and *ATM* (experimentally validated oncogenes that co-operate with mutant *KRAS* in tumorigenesis) and *EGFR* and *NF1* (both of which act in the RTK-RAS-RAF-ERK pathway). **C)** A similar analysis for non-silent mutations co-occurring with missense *EGFR* mutations, highlighting the negative association with *KRAS* mutations.

Several interpretations of such exclusivities have been proposed. When the mutated genes are thought likely to have similar consequences---for example, by encoding proteins that act in the same signaling pathway-a mutation in one could, in principle, obviate any selective advantage that might normally be conferred by the other. But there are more interesting possible explanations for exclusivity---a combination of mutations might result in synthetically lethal or at least growth-impairing phenotypes, by providing a toxic signal or impairing an essential cellular function.

Consider one example of a striking but still unexplained mutual exclusivity. Mutations affecting one of four different RNA splicing factors are commonly found in myelodysplasias and, less frequently, in certain solid tumors (Yoshida et al. 2011; Ogawa 2012; Lee 2016). These mutations change the transcriptome by favoring certain alternative choices during RNA processing, so they are fundamentally GOF mutations. Since the second (normal) copy of each gene is retained---and indeed cannot be eliminated without loss of viability (Fei et al. 2016)---the mutant version also appears to be deficient in some essential functions of the normal gene. But heterozygous mutations affecting two splicing factors have not been observed in a single tumor. This could be accounted for in either of two ways. Mutations affecting two factors might prevent a sufficient level of normal splicing. Alternatively, two mutant splicing factors might produce more aberrantly processed RNAs than growing cells could tolerate. In either view-the loss of normal function or a gain of aberrant function-effects on cell growth or viability could disadvantage a cell with two splicing factor mutations in comparison with other cells in the same cancer.

## Explaining a frequently encountered example of mutual exclusivity

We have been studying a common (and commonly noted) example of mutual exclusivity: the rarity of encountering co-existing *EGFR* and *KRAS* GOF mutations in human lung adenocarcinomas, despite the fact that each kind of mutation is found frequently (in 10 to 30 percent of tumors) without the other (Cancer Genome Atlas Research 2014)(Figure 1).

Interpretation of this observation has to be considered in the light of at least three important factors: the mutation rates in the relevant genes in a single cell (before, during, or after carcinogenesis)^1^; the effect of any mutations in that gene on the relative growth and survival rates of that cell; and the consequences of other coexisting mutation(s) that might confer a selective advantage or disadvantage on that cell. These features obviously complicate any analysis of tumor genotypes, since they can all affect the likelihood of detecting coincident mutations when DNA from a tumor sample is studied. It is equally important to consider the number of cells that exist in a tumor or in a tumorigenic state---from one to billions---when a mutation occurs in a single cell. This will dramatically influence the frequency with which mutations are detected in tumor tissue.

One way to approach the problem is to remove the bias of selective pressure from the analysis, examining tumors for the co-existence of a mutation known to be oncogenic and another mutation that does not alter coding potential. Assuming that the oncogenic mutation has little or no effect on the somatic mutation rates and that the non-coding mutation can provide no selective advantage or disadvantage whether it occurs with or without the oncogenic mutation, the frequency at which co-incident mutations are detected should be predictable from the frequencies at which either is detected alone: calculated simply as the product of the two mutation frequencies. Consistent with this idea, we have previously shown that such silent mutations are detected along with codon 12 KRAS mutations as the products of their individual frequencies (see Figure 1B in (Unni et al. 2015)).

But the situation is more complicated when we ask whether two mutations that change the coding sequence---and particularly two mutations that affect a cell’s selective advantage---are present in the same cell. If the mutation rate, starting cell number, and the selective advantages of either mutant gene are not affected by the presence of the other, the prediction is a simple one: co-existence of the two mutations should be detected at a frequency determined by multiplying the frequencies at which each mutation is detected singly. Thus if a mutation in gene A is found in 30 percent of tumors and a mutation in gene B is found in 10 percent, coincident detection of the two should then occur in about 3 percent of tumors. However, if the presence of the mutation in gene A eliminates any advantage conferred by the mutation in gene B or vice versa, as in the situation often referred to as “pathway redundancy,” then the likelihood of finding both mutations in the tumor will be markedly reduced. Conversely, if the co-existence the two mutations has a synergistic effect that exceeds the sum of the selective advantages conferred by each, then tumors with both mutations will be more common that predicted from the product of individual frequencies. Finally, and relevant to the problem we address here, if co-existence of mutations confers toxicity---a selective disadvantage---on cells, then (as in the loss of a selective advantage) tumors will less commonly---and perhaps never---be found to harbor both mutations.

This kind of analysis is germane to an observation made by several laboratories and studied mechanistically here: despite the relatively high frequencies of detection in human lung adenocarcinomas of codon 12 mutations in KRAS (about 30%) and a few idiosyncratic mutations in EGFR (about 10-15%), co-existence of the two is rarely, if ever, found (Ding et al. 2008; Cancer Genome Atlas Research 2014; Unni et al. 2015). We have assessed exon sequence data for over 500 lung adenocarcinomas, taking into consideration specific demographic factors associated with the incidence of these mutations such as smoking status, and we have found a significant negative association between the two mutant oncogenes.

An exome-wide survey for the co-occurrence of mutated genes with mutant KRAS in 520 lung adenocarcinomas revealed that mutations in EGFR deviated most dramatically from the frequency expected for situations in which mutation rates and selective advantages were unaltered by co-existence of the mutations (Figure 1). Importantly, the analysis also revealed mutated genes, such as *STK11*, that occur together with mutant KRAS more commonly than anticipated by chance. This combination has been posited to co-operate during tumor progression (Ji et al. 2007), suggesting that the method can reveal mutated gene pairs that occur more and less frequently than would have been predicted if co-occurrence did not influence selective advantages or mutation rates. For simplicity, the analysis did not take into account gene copy numbers or methylation changes; in principle, either could create GOF or LOF. Further, we have not considered the variables of time and allele frequency; the sequenced DNAs were obtained from tumors that had probably evolved for different numbers of cell divisions, and not all of the recorded mutations were necessarily in all the cells of the tumor.

In an initial effort to understand why mutations in *EGFR* and *KRAS* were not found in the same tumor at detectable frequencies, we took advantage of transgenic mice that we had engineered to express mutant alleles of human *KRAS* (G12D) and human *EGFR* (a small deletion in exon 19) in lung epithelium under the control of a doxycycline-responsive transcriptional regulator, rtTA (Fisher et al. 2001; Politi et al. 2006). We made the simple prediction that in tri-transgenic mice---carrying both oncogenic transgenes and a third encoding the regulator---tumors would develop more rapidly (and mice would be sacrificed earlier) if the oncoproteins had additive activities; tumors would appear at the same rate as in the mice with only the more potent transgene if there were no additive effects; and tumors would occur more slowly or not at all if the combination produced detrimental consequences.

What we observed, however, was more complicated and surprising. All mice developed multiple tumors throughout the lungs. As indicated by the Kaplan-Meier plots shown in Figure 2 in Unni et al (2015), the tumors affected the survival of the tri-transgenic animals at approximately the same rate as in the line of bi-transgenic animals having only the more potent of the two oncogenic transgenes (*KRAS*-G12D). This seemed to imply that addition of the mutant *EGFR* transgene had no further effect, neither providing a selective advantage nor impairing cell growth or viability in the presence of the *KRAS* transgene. However, when we tested individual microdissected tumors for expression of the two genes, RNA specific for only one of the two transgenes was found in the great majority of the tumors. The most likely explanation of this observation is that detectable tumors arose only when cells expressed a single oncogenic transgene, not both. The tumors we harvested and characterized presumably developed from cells that never expressed both transgenes or from cells that may have expressed both transgenes initially but then extinguished the expression of one of them. Since we do not know the proportion of lung epithelial cells that turn on each transgene immediately or soon after administration of doxycycline, it is difficult to know which scenario is more likely. Similar findings with other pairs of *EGFR* and *KRAS* transgenic mice have recently been published by (Ambrogio et al. 2016).

**Fig. 2.**
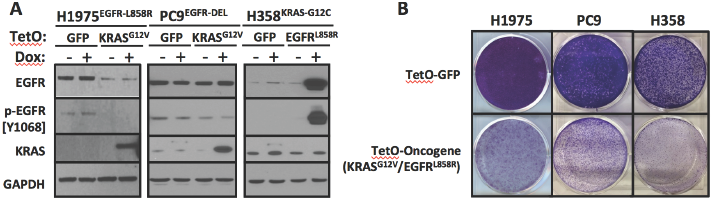
Co-expression of mutant KRAS and mutant EGFR in human lung adenocarcinoma cell lines decreases viability. **A)** Induced expression of transduced genes in human lung cancer cell lines. H1975, PC9 and H358 cell lines (with endogenous mutations in either *EGFR* or *KRAS* as indicated) were transduced with the indicated tetracycline (TetO) responsive plasmids. Lysates of cells cultured in the presence or absence of doxycycline (Dox) for 24 hours were prepared and assayed for the indicated protein expression by western blotting. **B)** Co-expression of mutant *KRAS* and *EGFR* reduces cell viability. Cells with either a TetO-Oncogene (*KRAS*^G12V^ for PC9 and H1975, *EGFR*^L858R^ for H358) or TetO-GFP were grown in the presence of Dox for seven days and cell density was assessed by crystal violet staining. Only the cells in which Dox induces expression of the second oncogene have reduced cell densities, consistent with the hypothesis that the co-expression of *KRAS* and *EGFR* leads to a less fit cellular state.

One of the shortcomings of the experiments with transgenic mice is that our claims for synthetic toxicity of the two expressed oncogenes is conjectural and not based on direct observation of noxious effects. Furthermore, the experiments were, of necessity, performed in mouse rather than human cells. To overcome these limitations, we placed inducible DNA vectors carrying a mutant *EGFR* or a mutant *KRAS* gene into human lung adenocarcinoma cell lines that express either of the two oncogenic drivers of the cancers from which the cell lines were derived. Thus PC9 cells, with an *EGFR*-exon 19 deletion mutation, were equipped with a doxycycline-inducible *KRAS* gene (G12D), and H358 cells, with a *KRAS*-G12C mutation, were modified by addition of an inducible *EGFR*-exon 19 mutation. PC9 and H358 lines carrying an inducible green fluorescent protein (GFP) cDNA served as controls. When the exogenous coding domains were induced in cultured cells, co-expression of a second oncogene (mutant *KRAS* in PC9 cells or mutant *EGFR* in H358 cells) caused a gradual decline in cell numbers, whereas expression of GFP produced no obvious changes. By 7 to 10 days after induction, only 10 to 20 percent as many lung cancer cells remained in the cultures in which a second oncogene was induced as in the control cultures, suggesting that cell death mechanisms were activated, with or without restriction of cell growth (Figure 2)

Several morphological, biochemical, and physiological properties were associated with the diminished numbers of cells in these cultures. Most obviously, many of the cells showed a marked change in shape and cytoplasmic contents. Electron micrographs revealed the kinds of vesicular structures associated with macropinocytosis, and the predicted augmented uptake of proteins was documented with fluorescent dextran (Unni et al. 2015). Since similar findings have been reported in cells with high levels of mutant KRAS proteins (Commisso et al. 2013), we also examined the abundance and phosphorylation status of several proteins implicated in transmission of signals downstream of KRAS and EGFR (Figure 3A). Of special note are the increased levels of phosphorylation of four serine/threonine kinases---AKT, ERK, p38, and JNK-suggesting that enzymatic activity in these branches of the RAS signaling network was augmented by coexpression of mutant EGFR and mutant KRAS (Figure 3B).

**Fig. 3.**
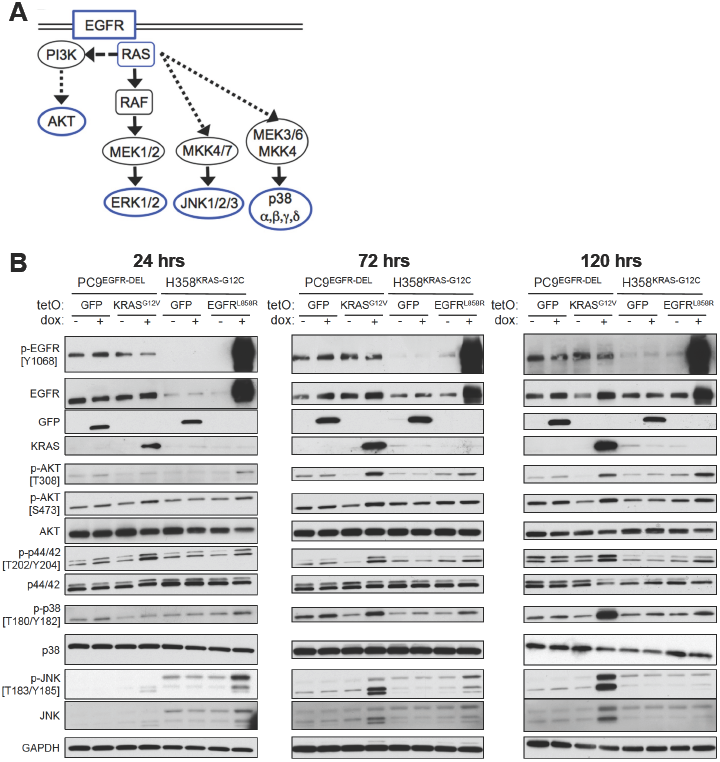
Downstream components of the EGFR-KRAS signaling axis are activated at different time-points after induction of a second mutant oncogene. **A)** EGFR and KRAS mainly mediate their oncogenic effects through induction of the PI3K-AKT and RAF-ERK pathways. However, KRAS has also been shown to activate additional MAPK signaling pathways including JNK and p38 under specific conditions, namely through association and activation of RAC (Coso et al. 1995; Minden et al. 1995; Bhanot et al. 2010). **B)** Temporal effects of mutant *KRAS* and *EGFR* co-expression on downstream signaling pathways over time. Increased MAPK signaling is observed in cells co-expressing mutant oncogenes; however, the times at which they are activated vary depending on the line and the specific MAPK pathway components (ERK, JNK or p38). The indicated cells were cultured for 24, 72 and 120 hrs with or without Dox; lysates were assayed by western blotting for the indicated proteins and phospho-proteins. Where relevant, the phosphorylated Tyrosine (Y) or Threonine (T) residue that determines reactivity with phospho-specific antibodies is shown.

When the tumor cell lines expressing both oncogenes were examined for markers of cell cycle progression, apoptosis, and autophagy, we could document perturbations of all these processes, and each could help to explain the loss of cell viability. Combined with the morphological changes attributed to increased cell vacuolization, we suggest that a unique form of cell death, termed “methuosis”, can be induced by co-expression of the mutant oncogenes (Maltese and Overmeyer 2014). However, whether this is true only for these experimental systems or is indeed a feature specifically associated with the synthetic lethality associated with mutant KRAS and EGFR remains to be explored.

One major factor that could influence the phenotype observed is the level at which the mutant oncogenes are expressed in the experimental systems. For example, our work utilized doxycycline inducible systems in mice and cell culture, and these can produce proteins at high levels (Unni et al. 2015). A recent study exploring the relationship of mutant *BRAF* and *KRAS* in lung adenocarcinoma, however, expressed the mutant genes from their endogenous promoters, and senescence rather than methuosis was observed upon co-expression (Cisowski et al. 2016). While the levels of the oncoproteins may influence the specific physiological effects, the premise that mutually exclusive mutant oncogenes lead to a less fit cellular state upon coexpression remains consistent.

## Searching for the mechanism and utility of agonistic synthetic lethality

To understand and exploit the apparent synthetic lethality resulting from coexpression of two oncogenes, it is necessary to identify the factors that mediate the toxic effects and to learn how to recapitulate them. We have considered and, in some instances, initiated attempts to do this in several ways.

i. An unbiased screen with reagents that mutate or inhibit the function of thousands of individual genes---through gene editing, inhibitory RNAs, or chemical compounds---might identify genes required for synthetic lethality. To that end, we have infected cultures of H358 cells carrying an uninduced *EGFR* (L858R) transgene, in addition to an endogenous somatic *KRAS* mutation, with a lentivirus library carrying CRISPR/Cas9 and guide RNAs targeting about 20,000 genes (Shalem et al. 2014). The infected cells were then treated with doxycycline to induce mutant *KRAS*, expecting that after a few passages the cultures would be enriched for cells in which gene editing had inactivated any gene required to mediate the toxic effects of oncogene co-expression. NGS sequencing of DNA from surviving cells revealed enrichment for cells in which CRISPR had inactivated *EGFR* (presumably the mutant transgene) or a variety of endogenous genes, including two genes encoding Mediator proteins.
ii. Enough is now known about the proteins involved in EGFR and KRAS signaling to make educated guesses about the components that might be critical for synthetic toxicity. A set of inhibitory RNAs has been developed to “knock-down” the relevant gene products individually to test whether they are required to impair the function of cells expressing both oncogenes. Furthermore, we have recently observed a dynamic relationship in the activation of MAPK pathway components upon coinduction of mutant *EGFR* and *KRAS* in different cell lines (Figure 3). This has led us to explore the kinetics of signaling upon co-induction of the mutant oncogenes using quantitative phospho-proteomics to identify the key components that mediate synthetic lethality
iii. Compounds that inhibit enzymes, such as protein kinases or phosphatases, that regulate the activity of components of EGFR and KRAS signaling networks could provide information about elements that contribute to---or protect cells from---the detrimental effects of EGFR plus KRAS on cell growth and survival. In principle, some compounds might be instructive about ways to use drug treatment of tumors to mimic the synthetic lethality produced by combinations of oncogenic mutations. Others might help to identify proteins that mediate the toxicity of the genetic combinations.

To attempt to understand synthetic agonistic lethality more generally, we encourage studies of recognized examples of mutually exclusive oncogenes and efforts to find oncogenic combinations that are detrimental to cell growth or survival. Among the several known oncogenic mutations that appear to be mutually exclusive, perhaps the best known, other than the *EGFR/KRAS* combination, is the rarity of an *NRAS* mutation coincident with a *BRAF* mutation in melanoma, despite the high frequency of each mutation in this tumor type. Further, Petti et al have reported that forced expression of mutant *NRAS* in a melanoma clone with mutant *BRAF* causes the cells to become senescent (Petti et al. 2006). We have surveyed TCGA databases for other missing combinations of mutant genes in many types of cancer, and some of those are listed in Table 1; direct tests of the effects of such combinations in appropriate cell types are likely to be informative. Another approach to identifying toxic combinations relies on the use of a library of genetic vectors carrying a large selection of activated oncogenes, preferably linked to bar codes. Propagation of cells into which such libraries have been introduced, combined with DNA sequencing to look for clones that become under-represented during growth, should produce candidate combinations for synthetic toxicity.

**Table 1.**
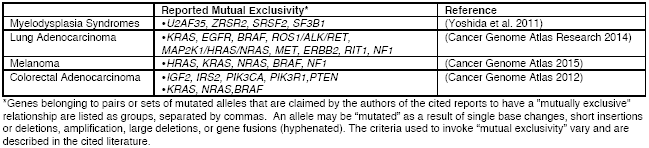

|  | **Reported Mutual Exclusivity\*** | **Reference** |
| --- | --- | --- |
| Myelodysplasia Syndromes | •*U2AF35, ZRSR2, SRSF2, SF3B1* | (Yoshida et al. 2011) |
| Lung Adenocarcinoma | •*KRAS, EGFR, BRAF, ROS1/ALK/RET, MAP2K1/HRAS/NRAS, MET, ERBB2, RIT1, NF1* | (Cancer Genome Atlas Research 2014) |
| Melanoma | •*HRAS, KRAS, NRAS, BRAF, NF1* | (Cancer Genome Atlas 2015) |
| Colorectal Adenocarcinoma | •*IGF2, IRS2, PIK3CA, PIK3R1, PTEN*<br>•*KRAS, NRAS, BRAF* | (Cancer Genome Atlas 2012) |

\*Genes belonging to pairs or sets of mutated alleles that are claimed by the authors of the cited reports to have a "mutually exclusive" relationship are listed as groups, separated by commas. An allele may be "mutated" as a result of single base changes, short insertions or deletions, amplification, large deletions, or gene fusions (hyphenated). The criteria used to invoke "mutual exclusivity" vary and are described in the cited literature.

A review of mutually exclusive oncogenic mutations shows that such pairs are not confined to GOF mutations in proto-oncogenes (Table 1). Several, such as mutations of *NF1* and *EGFR* in glioblastomas, involve a LOF mutation in a tumor suppressor gene plus an activated proto-oncogene. Of course, mutational loss of a tumor suppressor function may lead to a functional gain in a signaling protein that drives growth; examples might include the augmented signaling by a RAS protein after loss of a GAP (GTPase activator protein) or the increased HIF (hypoxia inducible factor) activity after loss of the Von Hippel-Lindau (VHL) protein. Further experiments to investigate why such combinations occur and to seek other detrimental pairings might enrich our understanding of the exclusivities and suggest additional strategies for therapies based on them.

## Summary: Interpretations and Further Prospects

Mutations in *EGFR* and *KRAS*, each found at high frequency in human lung adenocarcinomas, rarely if ever co-occur in a single tumor. Our efforts thus far to understand this phenomenon are most consistent with the notion that coexpression of the two mutations is synthetically lethal or at least toxic. It is, however, difficult to study the mechanism that explains the mutual exclusivity in an intact organism, even in mouse models, let alone human patients.

The most productive approach thus far has been to induce expression of the second oncogene in human adenocarcinoma cell lines that carry mutant *EGFR* or mutant *KRAS* as a driver oncogene after introduction of the second oncogene regulated by a doxycycline-responsive transcriptional activator. This provides a setting in which the effects of adding the second oncogene can be observed acutely, but it may not accurately recapitulate events in vivo, since the production of the two oncoproteins, especially the induced protein, may exceed levels encountered in the intact organism. Moreover, the growth conditions in tissue culture and the absence of the natural microenvironment may further alter the outcome. Thus it remains conceivable that the explanation for mutual exclusivity of certain oncogenic mutations in human cancers is a manifestation of other features. For example, tumor cells expressing both oncogenes might experience a growth disadvantage (e.g. senescence) without undergoing a lethal process. There might also be situations in which mutual exclusivity is, in fact, explained solely by functional redundancy of the relevant cancer genes, without a selective disadvantage conferred by the second gene contributing to the exclusivity.

Establishing alternative explanations for mutually exclusive pairs of oncogenic mutations will be difficult, and we have chosen to focus on efforts to understand the mechanisms by which cancer cells die in culture when the second oncogene is expressed. We have done this because the synthetic toxicity of *EGFR* and *KRAS* mutations is unexpected and its mechanism, even in cell culture, might be instructive about novel approaches to therapy. Many efforts have been made to screen for lethal outcomes in cancer cells (e.g. those in which common mutant *KRAS* alleles are oncogenic drivers) by using libraries of inhibitory RNAs, drugs that inhibit gene products, or CRISPR/Cas9-based libraries that inactivate genes (Downward 2015). In such screens, the intent is to combine a LOF event with a GOF mutation (e.g. in *KRAS)* and to cause cell toxicity by eliminating or severely reducing the functionality of a second gene that is specifically required for the viability of a cancer cell driven by a certain oncogene. Our findings imply that it might also be important to seek GOF changes---using agonists, not just antagonists, of a second gene’s function---to kill or disable cancer cells.

We refer here to GOF and LOF (or agonistic or antagonistic) approaches as a way of characterizing the action taken experimentally to perturb cancer cell behavior or viability. But the outcome may involve biochemical steps that are complex and difficult to classify as losses or gains of function. Cell signaling is still poorly understood, involving branching pathways, feedback events, and dosage. In this light, inhibition of a negative regulator (an apparent LOF) would have a positive effect on the regulated protein (a GOF). Further, by viewing events in cell signaling in a binary language---largely “on or off” or “activated or inactivated”-we may be ignoring important effects of variations in signal intensity (e.g. imperceptible when the signal is weak, oncogenic when moderate, and toxic when intense).

In addition to exploring the basis of the common mutual exclusivities, such as mutant *EGFR* and mutant *KRAS*, we encourage the further study of other combinations of oncogenic mutations that rarely or never co-occur. Those examples that have attracted attention include mutant *BRAF* and mutant *NRAS* (shown to cause cell senescence when forced to be expressed concurrently in melanoma cells (Petti et al. 2006)); the mutual exclusivity of splicing factor mutations; mutant *KRAS* and *BRAF* in lung adenocarcinoma (Cisowski et al. 2016); inactivation of *NF1* and activating *EGFR* insertions in glioblastoma; and mutant *KRAS* and mutant *BRAF* in colorectal adenocarcinoma (Table 1; (Yoshida et al. 2011; Cancer Genome Atlas 2012; Cancer Genome Atlas Research 2014; Cancer Genome Atlas 2015)). The list of mutually exclusive mutations, however, may be considerably longer, based on analysis of large data sets, such as those produced by TCGA. Since some genes in these analyses will have been infrequently mutated, it may be difficult to provide convincing evidence that sets of mutations are truly mutually exclusive, unless subjected to the kinds of experimental tests described here. Clues to the possible significance of exclusivity may be obtained by grouping genes by functional classes. Such approaches have been used, for example, to predict the mutual exclusivity of mutations affecting genes that modify chromatin in glioblastomas (e.g. *IDH1, MLL3, SETD1A;* (Brennan et al. 2013)) or affecting genes that encode protein kinases in acute myeloid leukemias (e.g. *FLT3, KIT, ABL1;* (Cancer Genome Atlas Research 2013)). Elucidating the mechanisms by which exclusivity is enforced in a variety of cancer types may suggest therapeutic strategies and enable chemical or genetic screens to pursue those strategies.

One other inadequately studied aspect of mutual exclusivity is the possible contribution of germ line variation to a cell’s ability to tolerate certain mutational combinations. Most of the evidence for inherited factors in carcinogenesis has come from the identification of germ line mutations in tumor suppressor genes, in genes that participate in DNA repair, and (least frequently) in proto-oncogenes. However, it seems likely that other kinds of genetic variation, responsible for less obvious changes in cancer risk, may occur at loci that determine the penetrance of oncogenic mutations in various cell lineages. Similarly, such variation may determine the probability of synthetic lethality due to combinations of GOF and LOF changes.

## Acknowledgements

The authors are grateful for support from the NIH Intramural Research Program, the Meyer Cancer Center at Weill Cornell Medicine, the British Columbia Cancer Foundation and the Canadian Institutes of Health Research (CIHR). W.L. also receives salary support from the CIHR New Investigator and Michael Smith Foundation for Health Research Scholar programs. We also thank our laboratory colleagues at the NIH, WCM, and UBC for helpful conversations.

## Footnotes

1 Mutation rates vary among genes, based on the size of the gene, its base composition, and its transcriptional activity; further, the average mutation rate in any cell will be affected by its exposure to DNA damaging agents, the fidelity of its replication machinery, and the integrity of its DNA repair mechanisms.

## Literature cited

Ambrogio C, Barbacid M, Santamaria D. 2016. In vivo oncogenic conflict triggered by co-existing KRAS and EGFR activating mutations in lung adenocarcinoma. Oncogene.

Baylin SB, Jones PA. 2016. Epigenetic Determinants of Cancer. Cold Spring Harb Perspect Biol 8.

Bhanot H, Young AM, Overmeyer JH, Maltese WA. 2010. Induction of nonapoptotic cell death by activated Ras requires inverse regulation of Rac1 and Arf6. Mol Cancer Res 8: 1358–1374.

Brennan CW, Verhaak RG, McKenna A, Campos B, Noushmehr H, Salama SR, Zheng S, Chakravarty D, Sanborn JZ, Berman SH et al. 2013. The somatic genomic landscape of glioblastoma. Cell 155: 462–477.

Cancer Genome Atlas N. 2012. Comprehensive molecular characterization of human colon and rectal cancer. Nature 487: 330–337.

Cancer Genome Atlas N. 2015. Genomic Classification of Cutaneous Melanoma. Cell 161: 1681–1696.

Cancer Genome Atlas Research N. 2013. Genomic and epigenomic landscapes of adult de novo acute myeloid leukemia. N EnglJMed 368: 2059–2074.

Cancer Genome Atlas Research N. 2014. Comprehensive molecular profiling of lung adenocarcinoma. Nature 511: 543–550.

Cerami E, Gao J, Dogrusoz U, Gross BE, Sumer SO, Aksoy BA, Jacobsen A, Byrne CJ, Heuer ML, Larsson E et al. 2012. The cBio cancer genomics portal: an open platform for exploring multidimensional cancer genomics data. Cancer Discov 2: 401–404.

Cisowski J, Sayin VI, Liu M, Karlsson C, Bergo MO. 2016. Oncogene-induced senescence underlies the mutual exclusive nature of oncogenic KRAS and BRAF. Oncogene 35: 1328–1333.

Commisso C, Davidson SM, Soydaner-Azeloglu RG, Parker SJ, Kamphorst JJ, Hackett S, Grabocka E, Nofal M, Drebin JA, Thompson CB et al. 2013. Macropinocytosis of protein is an amino acid supply route in Rastransformed cells. Nature 497: 633–637.

Coso OA, Chiariello M, Yu JC, Teramoto H, Crespo P, Xu N, Miki T, Gutkind JS. 1995. The small GTP-binding proteins Rac1 and Cdc42 regulate the activity of the JNK/SAPK signaling pathway. Cell 81: 1137–1146.

Ding L, Getz G, Wheeler DA, Mardis ER, McLellan MD, Cibulskis K, Sougnez C, Greulich H, Muzny DM, Morgan MB et al. 2008. Somatic mutations affect key pathways in lung adenocarcinoma. Nature 455: 1069–1075.

Downward J. 2015. RAS Synthetic Lethal Screens Revisited: Still Seeking the Elusive Prize? Clin Cancer Res 21: 1802–1809.

Fei DL, Motowski H, Chatrikhi R, Prasad S, Yu J, Gao S, Kielkopf CL, Bradley RK, Varmus H. 2016. Wild-Type U2AF1 Antagonizes the Splicing Program Characteristic of U2AF1-Mutant Tumors and Is Required for Cell Survival. PLoS Genet 12: e1006384.

Fisher GH, Wellen SL, Klimstra D, Lenczowski JM, Tichelaar JW, Lizak MJ, Whitsett JA, Koretsky A, Varmus HE. 2001. Induction and apoptotic regression of lung adenocarcinomas by regulation of a K-Ras transgene in the presence and absence of tumor suppressor genes. Genes Dev 15: 3249–3249.

Futreal PA, Coin L, Marshall M, Down T, Hubbard T, Wooster R, Rahman N, Stratton MR. 2004. A census of human cancer genes. Nat Rev Cancer 4: 177–183.

Garraway LA. 2013. Genomics-driven oncology: framework for an emerging paradigm. J Clin Oncol 31: 1806–1814.

Garraway LA, Lander ES. 2013. Lessons from the cancer genome. Cell 153: 17–37.

George J, Lim JS, Jang SJ, Cun Y, Ozretic L, Kong G, Leenders F, Lu X, Fernandez-Cuesta L, Bosco G et al. 2015. Comprehensive genomic profiles of small cell lung cancer. Nature 524: 47–53.

Hanahan D, Coussens LM. 2012. Accessories to the crime: functions of cells recruited to the tumor microenvironment. Cancer Cell 21: 309–322.

Hanahan D, Weinberg RA. 2000. The hallmarks of cancer. Cell 100: 57–70.

Hanahan D, Weinberg RA. 2011. Hallmarks of cancer: the next generation. Cell 144: 646–674.

Ji H, Ramsey MR, Hayes DN, Fan C, McNamara K, Kozlowski P, Torrice C, Wu MC, Shimamura T, Perera SA et al. 2007. LKB1 modulates lung cancer differentiation and metastasis. Nature 448: 807–810.

Lawrence MS, Stojanov P, Polak P, Kryukov GV, Cibulskis K, Sivachenko A, Carter SL, Stewart C, Mermel CH, Roberts SA et al. 2013. Mutational heterogeneity in cancer and the search for new cancer-associated genes. Nature 499: 214–218.

Lee SD, K.; Obeng, E. A.; Kim, E.; Micol, J. B.; Yoshimi, A.; Willekens, C.; Inoue, D.; Saada, V.; Cho, H.; Chung, Y. R.; Palacino, J.; Seiler, M.; Buonamici, S.; Smith, P. G.; Ebert, B. L.; Bradley, R.; Abdel-Wahab, O. 2016. Synthetic Lethal Interactions of MDS-Associated Spliceosomal Gene Mutations Identifies the Basis for Their Mutual Exclusivity. American Society for Hematology 58th Annual Meeting & Exposition: 961.

Maltese WA, Overmeyer JH. 2014. Methuosis: nonapoptotic cell death associated with vacuolization of macropinosome and endosome compartments. Am J Pathol 184: 1630–1642.

Meyerson M, Gabriel S, Getz G. 2010. Advances in understanding cancer genomes through second-generation sequencing. Nat Rev Genet 11: 685–696.

Minden A, Lin A, Claret FX, Abo A, Karin M. 1995. Selective activation of the JNK signaling cascade and c-Jun transcriptional activity by the small GTPases Rac and Cdc42Hs. Cell 81: 1147–1157.

Ogawa S. 2012. Splicing factor mutations in myelodysplasia. Int J Hematol 96: 438–442.

Petti C, Molla A, Vegetti C, Ferrone S, Anichini A, Sensi M. 2006. Coexpression of NRASQ61R and BRAFV600E in human melanoma cells activates senescence and increases susceptibility to cell-mediated cytotoxicity. Cancer Res 66: 6503–6511.

Politi K, Zakowski MF, Fan PD, Schonfeld EA, Pao W, Varmus HE. 2006. Lung adenocarcinomas induced in mice by mutant EGF receptors found in human lung cancers respond to a tyrosine kinase inhibitor or to down-regulation of the receptors. Genes Dev 20: 1496–1510.

Shalem O, Sanjana NE, Hartenian E, Shi X, Scott DA, Mikkelsen TS, Heckl D, Ebert BL, Root DE, Doench JG et al. 2014. Genome-scale CRISPR-Cas9 knockout screening in human cells. Science 343: 84–87.

Stratton MR, Campbell PJ, Futreal PA. 2009. The cancer genome. Nature 458: 719–724.

Unni AM, Lockwood WW, Zejnullahu K, Lee-Lin SQ, Varmus H. 2015. Evidence that synthetic lethality underlies the mutual exclusivity of oncogenic KRAS and EGFR mutations in lung adenocarcinoma. Elife 4: e06907.

Watson IR, Takahashi K, Futreal PA, Chin L. 2013. Emerging patterns of somatic mutations in cancer. Nat Rev Genet 14: 703–718.

Yoshida K, Sanada M, Shiraishi Y, Nowak D, Nagata Y, Yamamoto R, Sato Y, Sato-Otsubo A, Kon A, Nagasaki M et al. 2011. Frequent pathway mutations of splicing machinery in myelodysplasia. Nature 478: 64–69.

